# “Fluorescent minihelix assay for translation quantification in high-throughput cell-free systems”

**DOI:** 10.1101/2023.10.13.562088

**Authors:** Jessica A Willi, Michael C Jewett

## Abstract

We report the use of an optimized tetra-cysteine minihelix both as a fusion protein and as a standalone reporter in vitro with the Flash dye to study cell-free protein expression dynamics and engineered ribosome activity. The fluorescent product can be detected and quantified via its characteristic emission spectrum in a standard 96/384-well plate reader, RT-qPCR system, or gel electrophoresis. The fluorescent reporter helix is short enough to be encoded on a primer pair and can tag any protein of interest via PCR. Both tagged protein or standalone reporter can be detected in real time during and in terminal cell-free expression reactions, or in gel, without the need for staining. The fluorescent signal is stable and linearly correlates with protein concentration, thus is suitable for product quantification. Finally, we demonstrate how this reporter can be used for future efforts in engineering in vitro translation systems.

**Graphical abstract:** 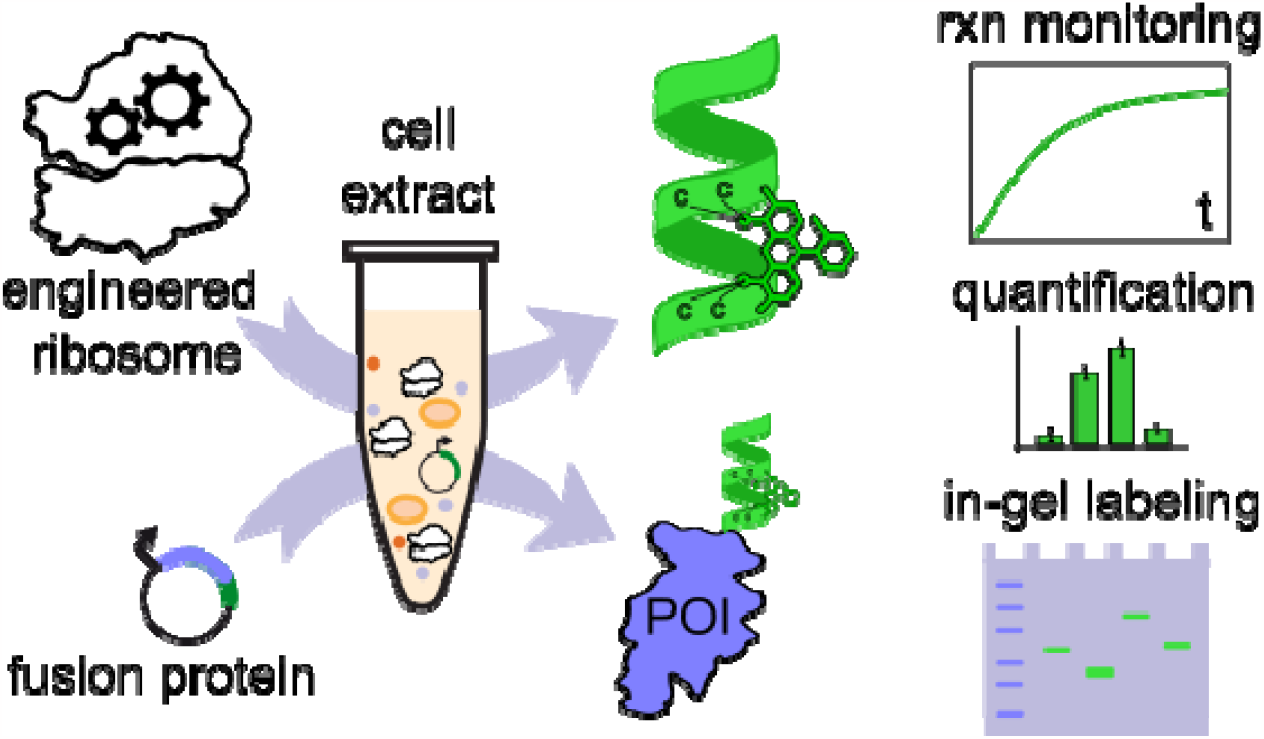

## Introduction

This assay is based on the pro-fluorescent dye 4,5-bis(1,3,2-dithiarsolan-2-yl)fluorescein (FlAsH-EDT_2_), discovered by the Tsien lab, which binds reversibly to a bipartite tetra-cysteine structure within two turns of an α-helix (Griffin et al., 1998). The dye molecule bound to the tetra-cysteine motif fluoresces in the green spectrum (absorption: 508 nm, emission 528-38 nm) The alpha helix was further optimized to allow for stronger binding of the FlAsH dye (Adams et al., 2002), and elevated quantum yield (Martin et al., 2005). The fluorescent protein tag/dye pair has been characterized *in vivo* for imaging and quantification (Adams et al., 2002; Adams & Tsien, 2008; Albert Griffin et al., 2000; Griffin et al., 1998; Hoffmann et al., 2010), and is even available as an in-cell detection kit (LumioGreen, Invitrogen), however, it has not been reported on for use within cell-free systems. Here, we express the optimized tetra cysteine helix both as a fusion protein and as a standalone reporter in an in vitro assay with the Flash dye to study cell-free protein expression dynamics and engineered ribosome activity.

## Results & Discussion

### Fetch assay for real-time reaction monitoring, protein quantification, and high-throughput assays

In a cell-free protein translation system (CFPS) based on E. coli extract, the reporter was transcribed from a T7 promoter-driven plasmid, producing an mRNA translated into the short fluorescence-emitting tetra cysteine helix (FETCH), whose sequence was previously optimized by (Martin et al., 2005). We found this assay to be highly flexible to be tailored along different characteristics (Fig. 1A): 1. The reporter minihelix can be either expressed as a standalone reporter of translation activity or as a tag to track the expression of protein of interest. 2. The FlAsH-EDT_2_ dye can be either be added to a completed cell free protein synthesis reaction (e.g. if reactions are split for other analysis), or at the start of the reaction, to obtain kinetic measurements without taking aliquots. 3. Just like CFPS, the assay scales well and reactions can be set up in microtubes, PCR tubes, or 96/384-well plates. 4. The fluorescent signal can be detected via plate reader, qPCR machine, or in-gel with UV-vis imaging. The flexible reaction and detection setup allows the FETCH to be used in many possible applications and in instrument-limited settings.

**Figure 1:**
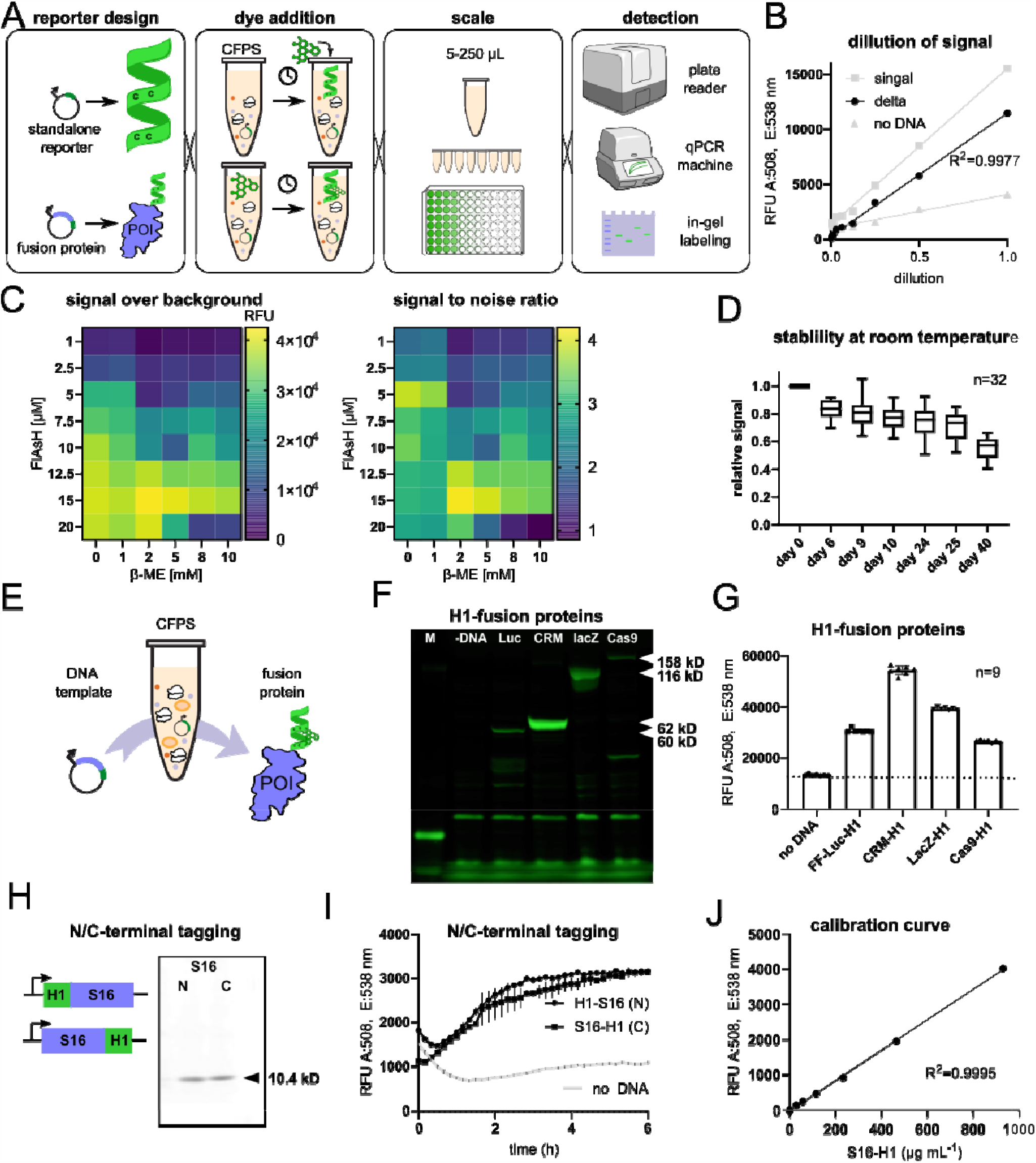
Optimization of FETCH in vitro assay for detection and quantification of CFPS products. A). The FETCH assay is flexible in reporter design, timing of the dye addition, scale and vessel format, and detection, all of which can be combined to tailor the assay to a multitude of applications. B). CFPS reactions expressing either H1 reporter helix (signal) or no DNA were serially diluted in PBS and their signal measured. The background-subtracted signal (delta) was fitted linear y and goes through 0/0 point. C). CFPS reactions expressing either H1 reporter or no DNA were substituted with varying concentrations of Flash-EDT2 dye and 2-mercaptoethanol, which competes with helix-dye binding and reduces background. Values were plotted for both RFU after background subtraction (left panel) or signal to noise (background) ratio (right). D). Plateaued CFPS reactions stored at room temperature on benchtop were measured over the course of weeks using the same instrument and program to determine total signal decay. Each well was normalized to its original value on day 0, whisker plot of n=32 samples. (E-G) Fusion proteins for tracking protein expression in CFPS. CFPS reactions expressing either FireflyLuciferase, CRM197-4xDQNAT, lacZ or Cas9 protein fused C-terminally were either F). separated on NuPAGE™ 4 to 12%, Bis-Tris in MOPS buffer and imaged with Azure gel imager on Cy3 setting or G). measured on a plate reader after 10h at 30*C. No DNA negative control represents the auto-fluorescence of the dye interacting with cysteine-rich proteins present in the extract. (H-J) Proteins can be tagged both N- or C-terminally: We tagged reporter H1 either up or downstream of ribosomal protein S16 from T. thermophilus and detected the CFPS-expressed fusion proteins either H). on 20% PAGE or I). in a plate reader. J). S16-H1 was expressed in the presence of ^14^C-Leucine and Flash-EDT, the reaction split into two equal serial dilutions, one of which was measured on a plate reader, the other was measured in a scintillation counter to determine the exact concentration of S16-H1 protein.

We found that the fluorescence signal after subtracting background is linearly proportional to the amount of reporter present (Fig. 1B), so it can be used for quantification purposes and scales with reaction volume. The presence of the FlAsH-EDT_2_ dye does not appear to interfere with transcription or synthesis. However, the dye can interact with other cysteine-rich proteins found in cell extracts or translation systems (Stroffekova et al., 2001), and produce a background signal in the absence of the desired tetra cysteine helix reporter. Thiol moieties such as EDT or ß-metcaptoethanol, or dithiotreol can quench this undesired background fluorescence. We found the final concentration of 15 μM FlAsH with 2 mM β-ME to be optimal signal to background for real-time observation of CFPS reactions (Fig. 1C). Due to the covalent bonds formed, the helix-dye product and resulting fluorescent signal are very stable. We stored plates of completed FETCH reactions at room temperature on the benchtop for multiple weeks, measuring every few days (Fig. 1D), and found that the fluorescent signal deteriorated at a rate of less than 3 % per day. This stability means that when live reaction monitoring is not required, researchers with limited plate reader capacity may set up many plates in parallel and quantify at the end point. This enables reaction setups in which thousands of ribosome variants or reaction conditions can be screened in a single experiment.

### Fetch fusion protein for tagging CFPS products

Fetch labelling offers a simple route to tag proteins produced cell-free synthesis whether detecting the protein of interest on a gel (an alternative to radiolabeling, FluoroTect, or western blotting), in real time (fusion to sfGFP) or quantifying final protein yield (split-GFP). For testing fusion proteins of various sizes, we C-terminally tagged firefly Luciferase, CRM197-4xDQNAT (a carrier protein used in conjugate vaccines), LacZ (beta-galactosidase, used in metabolic engineering and biosensors), and SpCas9 (used in genetic engineering applications). All fusion proteins were expressed and could be visualized in denaturing gel electrophoresis (Fig. 1F), where the fluorescent helix did not add significant mass to proteins, making each protein migrate at the expected kDa ranges. Since the dye is already covalently bound to the reporter helix, the gel can be imaged without staining after the run, even without removing it from the glass plates. Background binding can be reduced by addition of 25 mM 2-ME before loading the gel (Fig. S1A) and the specific signal allows for visualization of fusion protein products that would be indistinguishable from total lysate proteins in Coomassie staining (Fig. S1B). The same reactions quantified on a plate reader matched the expression levels of each protein (Fig. 1E). These setups also demonstrate the importance of running a no DNA control for background.

We found that the fluorescent helix can be added C- or N-terminally to proteins of interest, and its positioning did not appear to affect protein expression or the fluorescence yield in our example of *T. thermophilus* rspP (S16 protein) (Fig. 1H, I), although results may differ for proteins with highly structured ends. Real time reaction shape and slope correlates are affected by the tag position, protein length, complexity of the ORF: N-terminal emerge before C-terminal tags (Fig. 1I); smaller proteins emerge before longer, structured ORFs with rare codons. This means Fetch-tagging in CFPS presents a great tool for tracking translation dynamics of different ORFs (compare Fig. 1 I,E; Fig. 2 B, E). The plateau level is determined by the expression levels of the protein of interest or maximum available dye (which is typically not limiting with fusion proteins). Since fluorescence linearly correlates with protein concentration (Fig. 1B), Fetch can be used qualitatively: we created a standard curve by radiolabeling a Fetch-tagged rspP protein to determine its exact concentration in μg, then measured fluorescence emitted (Fig. 1J). We recommend creating a similar calibration curve in order to use Fetch to calculate μg yields for other proteins of interest.

**Figure 2:**
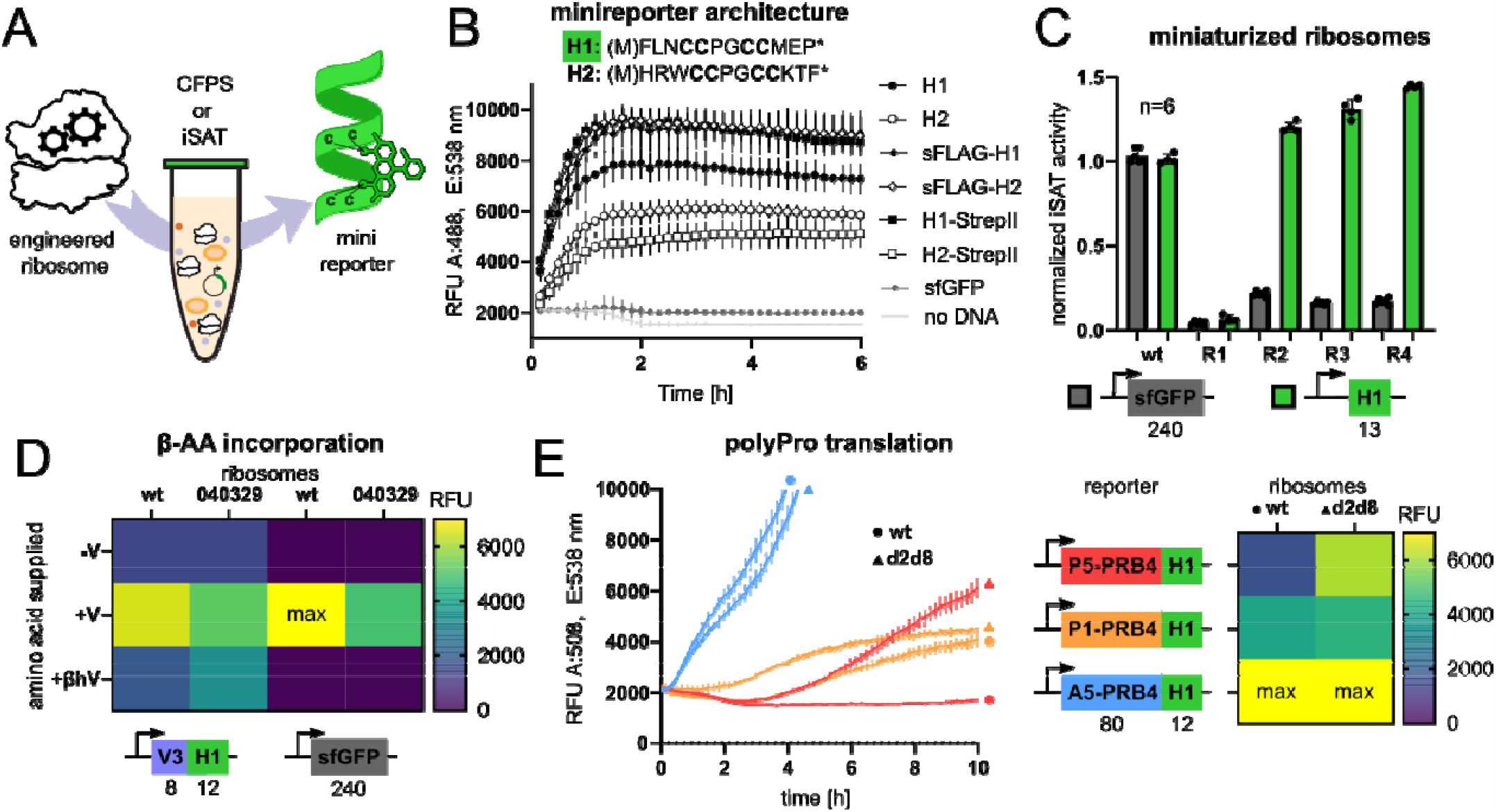
A). Fetch as standalone reporters for ribosome engineering. B). Real-time detection of reporter expression in plate reader. CFPS reactions expressing different reporter architectures, either standalone or fused to other functional tags sFLAG or StrepII tag. sfGFP is also expressed but emits low on FlAsH-specific channel. C). iSAT activity of ribosome constructs with engineered rRNA expressing either sfGFP or the H1-minireporter. Wt reactions are expressing a full-length 23S rRNA, while miniaturized ribosomes R1-R4 are expressing 23S rRNA variants with a 25nt deletion in domain II, replaced with different linkers. sfGFP and H1 were detected on their respective optimal channels, background was subtracted from all samples before normalizing to wt activity, bar graphs depict averages and standard deviation of n=6 experiments. D). Expression of Valine-coupled H1 reporter or sfGFP in the absence of valine, presence of valine, or β-homo-valine by either wt ribosomes or mutant 040329, which was evolved to accept β-AA better (Maini et al., 2015). Average RFU of n=3 reactions without background subtracted. E). iSAT expression of polyproline-rich protein PRB4 followed by H1 for detection, by either • wild type ribosomes or Δ d2d8 mutant, evolved specifically for increased polyproline expression (Schmied et al., 2018). Average RFU of n=3 reactions without background subtracted, heat map on the right depicts end point quantifications of polyproline reporters vs ribosomes.

### Fetch as a standalone reporter for ribosome engineering

When expressed as a standalone reporter, we found helix design H1 to yield higher signal than H2 (Fig. 2A), likely due to a more stable minihelix under CFPS conditions. H1 was also more tolerant of having functional sequences attached up- or downstream, such as purification tags sFLAG and StrepII, which indicates it is well suited for the construction of mini-reporter variants. Linkage to unstructured sequences can destabilize the minihelix, which can be remedied by including helix-forming MALEK residues in any additions. While the H1 sequence performed the best in our in vitro assay, the sequence can be altered to be shorter, contain other residues, or functional sequences both on the N or C-terminus of the helix (here StrepII, or sFLAG), as long as the central tetracysteine motif (CCPGCC) and helical structure is maintained (Fig. 2A). In order to use Fetch in ribosome engineering, we adapted it to iSAT, a system in which ribosomes are synthesized and assembled from a rDNA plasmid template, then functionally tested for translation capability (Fritz et al., 2015; Fritz & Jewett, 2014; Jewett et al., 2013). Traditionally the readout for iSAT is synthesis and fluorescence of sfGFP, however as we evolve and engineer ribosomes further from to the synthesis of novel polymers, they may also drift away from native translation, e.g., the synthesizing large α-amino acid-based proteins with the 20 canonical side residues. This functional drift has been observed in enzymatic evolution and is an acceptable payoff for novel catalysis. Therefore, requiring a readout of a large reporter protein is diametrically opposed to the development of alternative ribosomes with novel functions. We demonstrate that the Fetch assay is compatible with engineered ribosomes produced in iSAT (Supplementary Fig S2A), which allows for the high-throughput screening of rRNA variant designs.

We demonstrate that even translationally challenged ribosomes can more easily translate this short ORF compared to sfGFP. For instance, we used iSAT to construct a panel of ribosomes with rRNA containing deleted segment in domain II of 23S rRNA, and replaced them with different linkers (Fig. 2C, R1-4). While these mutated ribosomes had trouble expressing sfGFP (240 aa), some are markedly more successful at translating the mini-reporter of 13 aa. This suggests that the translational hurdle has been lowered in the Fetch assay, while still requiring the engineered ribosomes to engage in the hallmark steps of translation (initiation, elongation, release). The differential abilities of these mutant ribosomes would remain hidden if one were to only consider sfGFP expression as a benchmark of translation.

Fetch also presents an ideal reporter for quantifying in vitro non-canonical amino acid incorporation. Since the reporter helix does not natively contain Valine, we incorporated three Val codons via the sequence (M)LVMVLVM upstream of the reporter sequence, which did not hinder fluorescent readout, thanks to cushioning V with pro-helical residues. This short ORF allowed us to globally replace Valine with ß-h-Valine in this within cell-free system, which is charged onto tRNA^Val^ by the endogenous Val-RS. While wildtype ribosomes can produce some Fetch signal in the presence of ß-h-Val, ribosome mutant 040329, (Maini et al., 2015) is able to produce a stronger signal (Fig. 1D). Mutant 040329 was specifically evolved to better incorporate ß-amino acids, which in previous assays were introduced via custom orthogonal tRNA synthetases or flexizyme-charged orthogonal tRNAs (Dedkova et al., 2012; Dedkova & Hecht, 2019; Maini et al., 2015). In contrast, this assay leverages the mini-reporter to produce a high-throughput compatible signal despite global replacement of amino acids with non-canonical alternatives, circumventing the need for orthogonal translation machinery. The same is not possible with GFP (Fig. 1D), in which neither the wild type nor the 040329 ribosome produce a signal in the presence of ß-h-Val.

Fetch can be combined with challenging ORFs to track ribosome evolution towards specific sequence specialization. For instance, stretches of polyproline are challenging for native ribosomes to translate, due to proline’s backbone clashing with rRNA residues, which induces stalling and requires resolution by EF-P (Huter et al., 2017). To measure polyproline stalling, we constructed a reporter containing 5 prolines, a segment of polyproline-rich protein PBR4, and reporter helix H1 (Fig. 2E, P5). Other reporters contained the same segments but only one proline (P1) prior to PBR4; and a proline-free control with the identical sequence with every proline replaced with alanine (A5), in both the pre-sequence and PBR4. The expression of all constructs was tracked via the florescence of downstream minihelix H1. Indeed, wild type ribosomes could only efficiently express A5 reporter, while engineered ribosome d2d8 was able to synthesize polyPro stretches in both the P5 and P1 constructs. Ribosome mutant d2d8 is the products of previous directed evolution towards specialization in polyPro translation (Schmied et al., 2018). While this assay is also possible with GFP instead of FETCH, it is likely that proline-friendly ribosome variants would be less processive in translating proline-poor sequences such and would thus have a disadvantage synthesizing GFP compared to wt ribosomes. The fluorescent minihelix provides a high-throughput friendly signal without placing a significant translational burden on engineered ribosomes.

We propose that this assay may enable new avenues to evolve the ribosome towards minimized rRNA, non-canonical amino acids, or challenging open reading frames, by providing a fluorescent signal with a minimal translational burden for modified ribosomes.

**Supplementary Figure S1:**
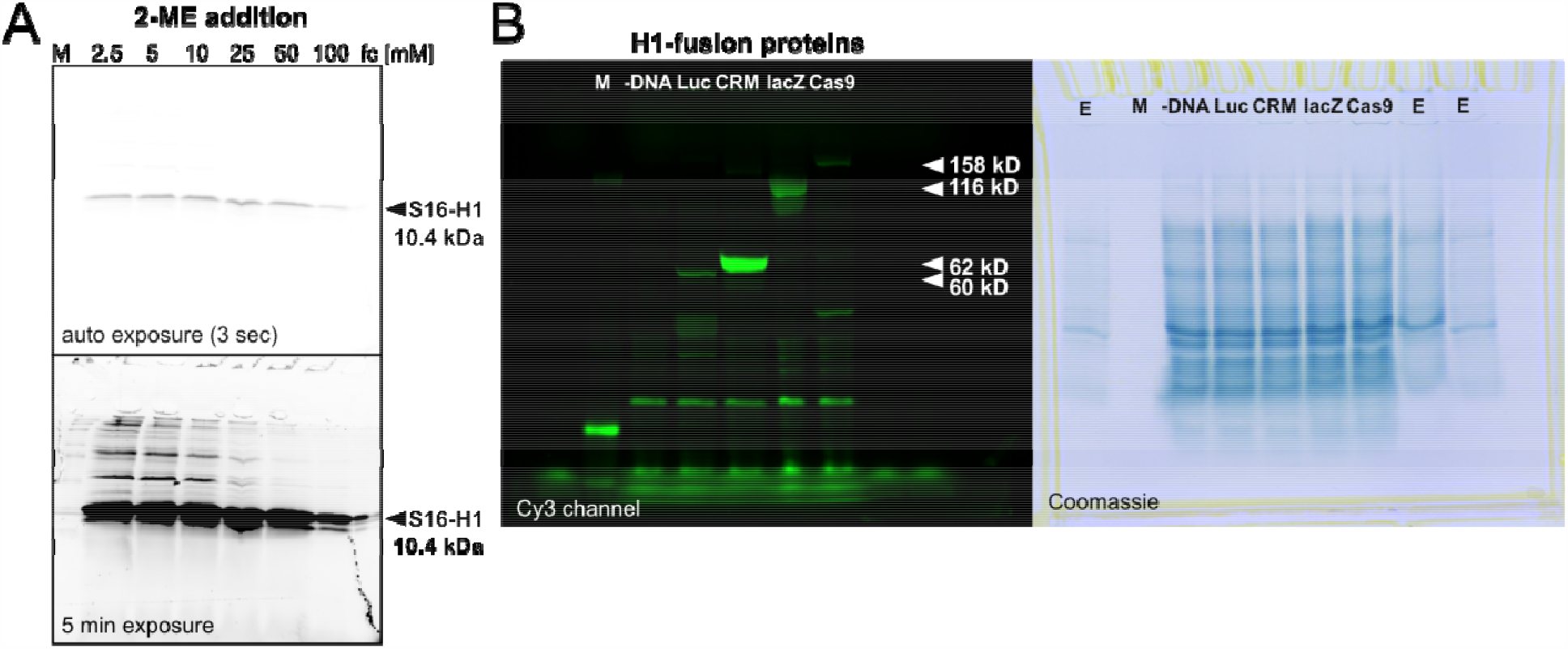
Imaging fusion proteins in gel. A). 2-ME addition after incubation with dye but before gel loading can reduce background fluorescence of FlAsH dye. Completed 5 uL CFPS reactions expressing S16-H1 fusion protein already containing Flash dye were brought to 15 uL with PBS, mixed with 5 uL 4x NuPAGE LDS Sample Buffer (Thermo Fisher) and heated to 70*C for 10 min. Indicated concentrations of 2-mercaptoethanol were added to the samples before loading onto 20% Bis-Tris in Tris-glycine buffer to separate. Image was captured on cytiva Cy3 channel on either auto-exposure (top, 3s) or overexposed (bottom, 5min) to show background fluorescence and Invitrogen BenchMark Fluorescent Protein Standard (left, M) B). Uncropped image of Cy3 channel and Comassie stain of gel shown in Fig. 1F, total proteins of spent CFPS reaction separated in NuPAGE™ 4 to 12%, Bis-Tris in MOPS buffer. Diluted total protein was also loaded on adjacent empty wells to limit bleeding (sides, E).

**Supplementary Figure S2:**
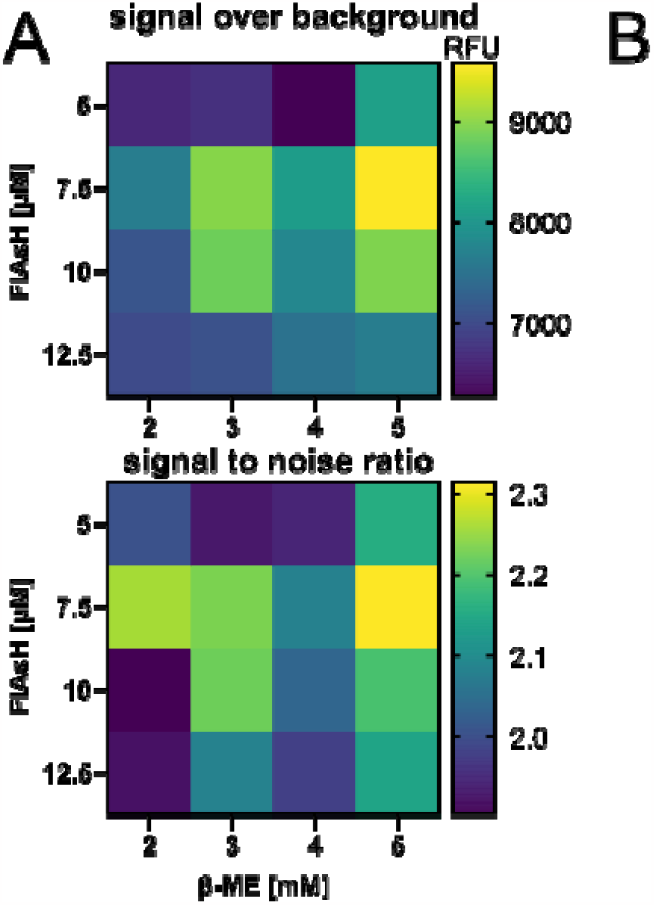
Optimization of FETCH assay for use in iSAT (in vitro ribosome synthesis, assembly, and translation). A). CFPS reactions expressing either sFLAG-H1 reporter or no DNA were substituted with varying concentrations of Flash-EDT_2_ dye and 2-mercaptoethanol, which competes with helix-dye binding and reduces background. Values were plotted for both RFU after background subtraction (upper panel) or signal to noise (background) ratio (bottom). B). Placeholder for further Supplementary Figure

## Materials and Methods

### Dye preparation

FlAsH-EDT_2_ (Cayman Chem, # 20704) was suspended in DMSO to a concentration of 500 μM, stored at -20°C. A fresh 15x concentrated mixture of FlAsH-EDT_2_ and 2-Mercaptoethanol (Sigma-Aldrich) was prepared in PBS buffer and incubated for 15 min at RT, prior to adding to the CFPS reaction. Final concentrations of the dye were optimized for each use case and are detailed below.

### Reporter plasmids

Fetch reporters were constructed on a pJL1 plasmid (T7 promoter, KanR) compatible with our protein synthesis assays. All reporters were constructed by site-directed mutagenesis via PCR sing Q5® High-Fidelity DNA Polymerase (NEB) to create overlaps forming the Fetch sequence followed by Gibson assembly and transformation into DH10β cells. The Fetch variants (H1: FLNCCPGCCMEP and H2: HRWCCPGCCKTF) were either cloned directly between start and stop codon of the open reading frame of pJL1, or added either downstream of sFLAG tag, or upstream of StrepII tag. H1 was also cloned C-terminally to pJL1-CRM197-4xDQNAT, -LacZ, and -SpCas9, as well as V3, polyPro reporters P5, P1 and A5. All reporters were confirmed by Sanger sequencing and deposited to Addgene (see Supporting Information Addgene #). Plasmid pJL1-sfGFP (Addgene #102634) was used as a control for CFPS activity. All plasmids were isolated from cells via miniprep (Zymo kit).

### CFPS reactions

Cell-free protein synthesis reactions were prepared using BL21 Star (DE3) cell lysate (Silverman et al., 2019). 5 uL CFPS reactions were set up as described before, including 0.1 μg/μL T7 pol, 2 mM DTT, 13 ng/μL reporter plasmid, and the optimized final concentration of 15 μM FlAsH-EDT_2_ and 2 mM 2-ME unless otherwise specified (in the case of real time monitoring). CFPS reactions were incubated at 30*C for 10h while monitoring as described below. Alternatively, dye-free reactions can be set up and incubated for 30*C for 20h to completion, then mixed 1:1 with 10 μM FlAsH-EDT_2_ and 10 mM 2-ME in PBS and incubated at room temperature for 1h before quantification.

Detection of tetracysteine-FlAsH complex was ideally performed in a Biotek Synergy H1 plate reader by defining a read channel specific to FlAsH (absorption 508 nm, emission 538 nm) or alternatively in a Biorad CFX96 Touch Real-Time PCR Detection System by using the “all channels” setting and later selecting the HEX channel (Ex 515-535, Em 560-580) to evaluate. A negative control reaction containing no reporter DNA was always run to determine background fluorescence, specifically dye binding to cell lysate. A reaction expressing sfGFP can serve as a positive control for extract activity but should be monitored on a separate channel (absorption 485 nm, emission 528 nm). End points values were quantified by averaging the last 5 data points of a given reaction.

### Radiolabeling

Total CFPS yields of T. thermophilus S16 protein were quantified by incorporation of ^14^C-Leucine (PerkinElmer) as previously described (Kim et al., 1996). ^14^C-Leucine was included in CFPS reactions to reach a final concentration of 10 μM in triplicate 15 μL reactions and incubated overnight at 30 °C. Five μL of each reaction was mixed with an equivalent volume of 0.5N KOH and incubated for 20 minutes at 37 °C to deacylated tRNA. Five μL of each reaction mixture was then spotted onto two separate 96-well filter mats (PerkinElmer 1450-421) and dried under a heat lamp. One of the mats was washed three times for 15 min in 5% TCA at 4 °C to precipitate protein, and washed once in 100% EtOH before being fully dried under a heat lamp. Radioactivity was measured by a liquid scintillation counter (PerkinElmer MicroBeta) compared to the unwashed filter mat. Protein concentration was calculated from the measured scintillation count per reaction volume by total number of Leu per S16 protein (7L), ^14^C-Leu specific activity (300mCi/mmol), and molar mass of S16 (10.4 kDa).

### Gel separation and imaging

5 uL CFPS reactions already containing Flash dye were brought to 15 uL with PBS, mixed with 5 uL 4x NuPAGE LDS Sample Buffer (Thermo Fisher) and heated to 70*C for 10 min. 25 mM 2-mercaptoethanol was added to the samples before loading to further reduce background. Proteins were separated on appropriate polyacrylamide gels, eg NuPAGE™ 4 to 12%, Bis-Tris in MOPS running buffer for larger proteins, and 20% Bis-Tris in Tris-glycine buffer for products below 40 kDa. Gels were run at 100 V with BenchMark™ Fluorescent Protein Standard (Thermo) and imaged immediately after completion using either Cytiva Amersham ImageQuant™ 800 or Azure Biosystems 200 imager on channel Cy3 (535 nm/UV). Coomassie staining of total proteins was performed using InstantBlue (Novus Bio) for gel documentation purposes.

### iSAT reactions

iSAT reactions allow for testing ribosomal variants encoded in a plasmid in a cell free translation system, as previously described (ref). S150 extract and ribosomal proteins (TP70) were prepared as previously described (ref), and reactions set up as detailed therein, with each reaction containing 3.1 ng/uL pJL1-reporter plasmid as specified in each Figure and 14 ng/uL pT7rrn plasmid encoding the ribosomal RNA variants, as well as 7.5 μM FlAsH-EDT_2_ and 5 mM 2-ME. Reactions were incubated at 37*C for 10h and fluorescence detected as described above.

